# Molecular markers characterization determining cell fate specification in an adult neurogenesis model of *Alzheimer’s disease*

**DOI:** 10.1101/2020.08.06.239111

**Authors:** Idoia Blanco-Luquin, Juan Cabello, Amaya Urdánoz-Casado, Blanca Acha, Eva Ma Gómez-Orte, Miren Roldan, Diego R. Pérez-Rodríguez, Maite Mendioroz

## Abstract

Adult hippocampal neurogenesis (AHN) study is still a challenge. In addition to methodological difficulties is the controversy of results derived of human or animal system approaches. In view of the proven link between AHN and learning and memory impairment, we generated a straightforward *in vitro* model to recapitulate adult neurogenesis in the context of Alzheimer’s disease (AD).

Neural progenitor cells (NPCs) monolayer culture was differentiated for a period of 29 days and Aβ peptide 1-42 was administered once a week. mRNA expression of *NEUROD1, NCAM1, TUBB3, RBFOX3, CALB1* and *GFAP* genes was determined by RT-qPCR.

Phenotypic changes were observed during directed differentiation. Except for *GFAP* and *CALB1*, these changes correlated with altered expression profile of all genes since 9 days. Only *TUBB3* expression remained constant while *NEUROD1, NCAM1* and *RBFOX3* expression increased over time. Moreover, Aβ treated NPCs showed transient decreases of mRNA expression for *NCAM1, TUBB3* and *RBFOX3* genes at 9 or 19 days.

Our *in vitro* human NPCs model is framed within the multistep process of AHN in the SGZ of the DG. Remarkably, its transcriptional assessment might reflect alterations detected in AD human patients, deepening our understanding of the disorder and possibly of its pathogenesis.

**SUMMARY STATEMENT:** Transcriptional profile of a number of genes recapitulating particular stages of Adult hippocampal neurogenesis in the context of Alzheimer’s disease

## INTRODUCTION

Adult neurogenesis is the process of forming functional neurons *de novo*. In an adult mammalian brain, neurogenesis occurs predominantly in specific brain niches: the subgranular zone (SGZ) of the dentate gyrus (DG) of the hippocampus and the subventricular zone (SVZ) lining the lateral ventricles (Hollands, Bartolotti and Lazarov, 2016; Hsieh, 2012). During the process of adult hippocampal neurogenesis (AHN), neural stem cells (NSCs) self-renew and differentiate, giving rise to transient amplifying progenitors (TAPs), neuroblasts, and eventually mature neurons, astrocytes, and oligodendrocytes. AHN regulators can be divided into intrinsic or extrinsic factors, that is, transcription factors (TFs) synthesized by the developing neural precursors and neurons, and growth factors and neurotrophins secreted from the surrounding niche, respectively (Covic, Karaca and Lie, 2010).

Understanding of human nervous system biology relies on the availability of human NSCs and their differentiated derivatives. A great deal of effort has been put into the generation of neural progenitor cells (NPCs) and neurons *in vitro*, mostly by differentiation of pluripotent stem cells (PSCs) and more recently by genetically reprogramming somatic cells to an embryonic stem (ES) cell-like state (induced pluripotent stem cells, iPSCs) (Yu, Marchetto and Gage, 2014). NPCs constitute an intermediate stage between pluripotent immature cells, such as ES cells and mature differentiated neural cells.

AHN clearly emerges as a robust phenomenon during both physiological and pathological aging in humans (Mu and Gage, 2011). In addition, the fact that it is implicated in normal functionality of hippocampal circuits, demonstrates an important link between adult neurogenesis and cognitive processes (Baglietto-Vargas *et al.*, 2017). As a consequence, impaired neurogenesis may negatively impact the survival of adult-born neurons and contribute to learning and memory failure, such as the associated with aging and disorders such as Alzheimer’s Disease (AD) (Mu and Gage, 2011; Coronel *et al.*, 2018; Li, Bao and Wang, 2016).

AD is the most common neurodegenerative disorder, characterized by progressive memory loss and cognitive decline caused by widespread neurons loss and synaptic connections in the cortex, hippocampus, amygdala and basal forebrain, and by a gradually significant loss of brain mass. Amyloid precursor protein (APP) plays a key role in normal brain development by influencing NSC proliferation, cell fate specification and neuronal maturation (Coronel *et al.*, 2018). However, its derivative Amyloid β (Aβ) peptide, a cleavage product of APP enzymatic processing, is the major component of the amyloid plaques, one of the hallmark pathologies found in brains of late-onset sporadic AD patients. Monomeric Aβ can self-aggregate to form oligomers, protofibrils, and amyloid fibrils which deposit as these amyloid plaques. Despite Aβ impact on neurogenesis is still controversial, Aβ plaques can cause severe damage to neurons and astrocytes, resulting in a gradual loss of neurons behind AD symptoms (Li, Bao and Wang, 2016).

Interestingly, noteworthy alterations in AHN have been detected at early stages of the disease, even before the onset of hallmark lesions or neuronal loss (Mu and Gage, 2011; Moreno-Jimenez *et al.*, 2019). Hence, a better understanding of AHN impairment observed at initial and later stages of AD by noninvasive methods might reveal insights into the pathogenesis of AD. What is more, restoration of normal levels of AHN would provide a potential therapeutic strategy to delay or halt AD-linked cognitive decline (Mu and Gage, 2011; Moreno-Jimenez *et al.*, 2019).

Here, we propose an intuitive *in vitro* approach to follow a stepwise lineage progression, as occurs during *in vivo* neurogenesis, by using human NPCs derived from an iPSC line as the starting source material. In order to find out whether human NPCs differentiation into mature neurons of any functional classification is disrupted in the AD microenvironment, we generated an *in vitro* model triggered by prolonged exposure towards nanomolar concentrations of Aβ peptide and characterized it by assessing a set of neurogenesis markers.

## MATERIALS AND METHODS

### NPCs culture, neuronal differentiation and Aβ peptide administration

Neural Progenitor Cells (NPCs) Derived from XCL1 DCXpGFP (ATCC® ACS5005™) were cultured following manufacturer recommendations. Briefly, 0.30 x10^6^ NPCs were seeded onto a CellMatrix Basement Membrane Gel (ATCC® ACS30235™) coated 12-well plate and incubated in NPC expansion medium: complete growth medium including DMEM/F-12 (Gibco, Fisher Scientific) supplemented with the Growth Kit for Neural Progenitor Cell Expansion (ATCC® ACS3003) and then maintained in a humidified incubator (5% CO2, 37 °C).

Neuronal differentiation experiments were carried out for 9, 19 and 29 days by plating NPCs at a seeding density of 80,000 viable cells/cm2 in 6-well coated culture plates. First, NPCs were incubated in expansion medium, which is referred to as day 0 in this study. Day post seeding, half-medium changes with differentiation medium were performed every 2-3 days throughout the duration of the culture period. Complete Differentiation Medium consisted of serum-free neuronal basal BrainPhys™ Neuronal Medium formulated to improve the electrophysiological and synaptic properties of neurons (Satir *et al.*, 2020), NeuroCult™ SM1 Neuronal Supplement (1:50), N2 Supplement-A (1:100), Recombinant Human Brain-Derived Neurotrophic Factor (BDNF, 20 ng/mL), Recombinant Human Glial-Derived Neurotrophic Factor (GDNF, 20 ng/mL), Dibutyryl-cAMP (1 mM) and ascorbic acid (200 nM) (STEMCELL Technologies, Vancouver, BC, Canada). Half fresh medium containing Amyloid β Protein Fragment 1-42 (50 nM; Sigma-Aldrich, St. Louis, MO, USA) or DMSO (Sigma-Aldrich) as vehicle was added once a week.

NPCs were harvested at 0 or 9, 19 and 29 days of differentiation for both conditions, by detaching with Accutase (Innovative Cell Technologies, San Diego, CA), washed with Dulbecco’s phosphate-buffered saline (DPBS, Sigma-Aldrich), centrifuged at 13000 r.p.m. and frozen at −80 ºC. All experiments were performed in triplicate.

### Neurogenesis markers mRNA expression analysis by RT-qPCR

Total RNA was extracted from frozen cell pellets of basal NPCs and control or Aβ treated NPCs incubated in differentiation media for 9, 19 or 29 days using RNeasy Mini kit (QIAGEN, Redwood City, CA, USA), following manufacturer’s instructions. Genomic DNA was digested with DNase I (RNase-Free DNase Set, Qiagen). Concentration and purity of RNA were both evaluated with NanoDrop spectrophotometer. Only RNA samples showing a minimum quality index (260 nm/280 nm absorbance ratios between 1.8 and 2.2 and 260 nm/230 nm absorbance ratios higher than 1.8) were included in the study. Complementary DNA (cDNA) was reverse transcribed from 1000 ng total RNA with Superscript® III First-Strand Synthesis Reverse Transcriptase (Invitrogen, Carlsbad, CA, USA) after priming with oligo-d (T) and random primers. RT-qPCR reactions were performed in duplicate with Power SYBR Green PCR Master Mix (Invitrogen, Carlsbad, CA, USA) in a QuantStudio 12K Flex Real-Time PCR System (Applied Biosystems, Foster City, CA, USA). Sequences of primer pair were designed using Real Time PCR tool (IDT, Coralville, IA, USA) and are listed in Supplementary Table S1. Relative expression mRNA levels of lineage specific genes in a particular sample were calculated as previously described (Livak and Schmittgen, 2001) and geometric mean of *ACTB* and *GAPDH* genes was used as the reference to normalize expression values.

### Immunofluorescence Staining

NPCs were seeded on Nunc™ Lab-Tek™ II chamber slides (Thermo Fisher Scientific, Waltham, MA, USA), coated with CellMatrix Basement Membrane Gel. Cells were either left untreated, or treated with Amyloid β Protein Fragment 1-42 (50 nM), in differentiation media as described above, and after 9, 19 or 29 days of incubation, fixated in 4% formalin (OPPAC, Spain) for 15 min. Cells were permeabilized using 0.5% TWEEN® 20 (Sigma-Aldrich) in DPBS, then blocked with 10% fetal bovine serum (Sigma-Aldrich) containing 0.5% Tween in DPBS 30 min at room temperature. Rabbit monoclonal anti-NeuN [EPR12763] (Cat# ab177487, RRID:AB_2532109; 1:300), anti-GFAP [EP672Y] (Cat# ab33922, RRID:AB_732571; 1:300), anti-Synaptophysin [YE269] (Cat# ab32127, RRID:AB_2286949; 1:200) and anti-Ki67 [SP6] (Cat# ab16667, RRID:AB_302459; 1:500) primary antibodies (Abcam, Cambridge, UK) diluted in blocking buffer were added, and incubated overnight at 4 ºC. After three washing steps, Alexa Fluor® 647 donkey anti-rabbit secondary antibody (Abcam Cat# ab150075, RRID:AB_2752244; 1:500) was added, and incubated for 30 minutes at room temperature in the dark. Following three washing steps, the slides were mounted with ProLong™ Gold Antifade Mountant with DAPI (Molecular Probes, OR, USA). Immunofluorescence images were acquired using a Cytation 5 Cell Imaging Multi-Mode Reader and analyzed with Gen5™ software (BioTek, Winooski, VT, USA).

### Statistical data analysis

Statistical analysis was performed with SPSS 21.0 (IBM, Inc., USA) and GraphPad Prism version 6.00 for Windows (GraphPad Software, La Jolla, CA, USA). First, we checked that all continuous variables showed a normal distribution, as per one-sample Shapiro-Wilk test. Data represents the mean ± standard error of the mean (SEM). Significance level was set at p-value < 0.05. Statistical significance for neuronal lineage specific genes mRNA levels across different time points was assessed by one-way analysis of variance (ANOVA) followed by *post-hoc* analyses Tukey’s honestly significant difference (HSD) and Dunnett’s multiple comparisons tests. In cases where Levene test did not show homogeneity of variance, Welch’s ANOVA followed by Dunnett’s T3 was conducted.

Paired t-test was used to analyze differences in the expression levels of the studied genes mRNA between Aβ treated and control group at each time point. GraphPad Prism version 6.00 for Windows was used to draw graphs.

## RESULTS

### Time-related changes in cultured NPCs during neural differentiation

To determine whether neural differentiation was effectively induced, we observed the morphological modifications of the cells over time. As shown in Figure 1, NPCs exposure to differentiation medium caused an increase in the number and length of the neuritic extensions, which can even connect with the extensions of neighbor cells, compared with basal cells grown in proliferation medium at zero time. These changes in cell morphology are typical of cells undergoing differentiation (Compagnucci *et al.*, 2016), were noticed since the first time point at 9 days, and became more evident as time went on in response to directed neurogenesis. Furthermore, total cell number in NPCs cultures remained steady, due to an absence of proliferation, stated by unchanged Ki 67 protein marker expression (Supplementary Figure S1), which correlated with a gradually cell differentiation boost. As a matter of fact, immunofluorescence (IF) staining proved NeuN (neuronal nuclei), GFAP (glial fibrillary acidic protein) and Synaptophysin protein expression, which mark neurons, glial cells and synaptic vesicles respectively, across the NPCs culture (Figure 2).

**Figure 1.**
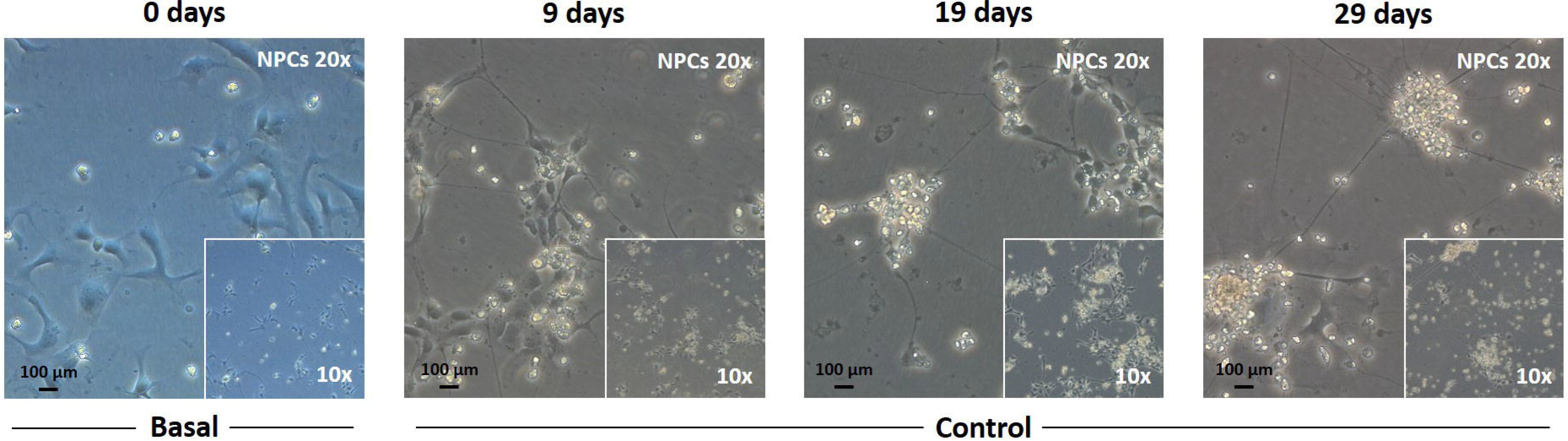
Phenotypic examination of NPCs directed differentiation in culture. Phase-contrast images at 0, 9, 19 and 29 days of basal cells incubated in expansion medium and control cells incubated in differentiation medium (20 x magnification with 10 x magnification inset, scale bar 100 μm).

**Figure 2.**
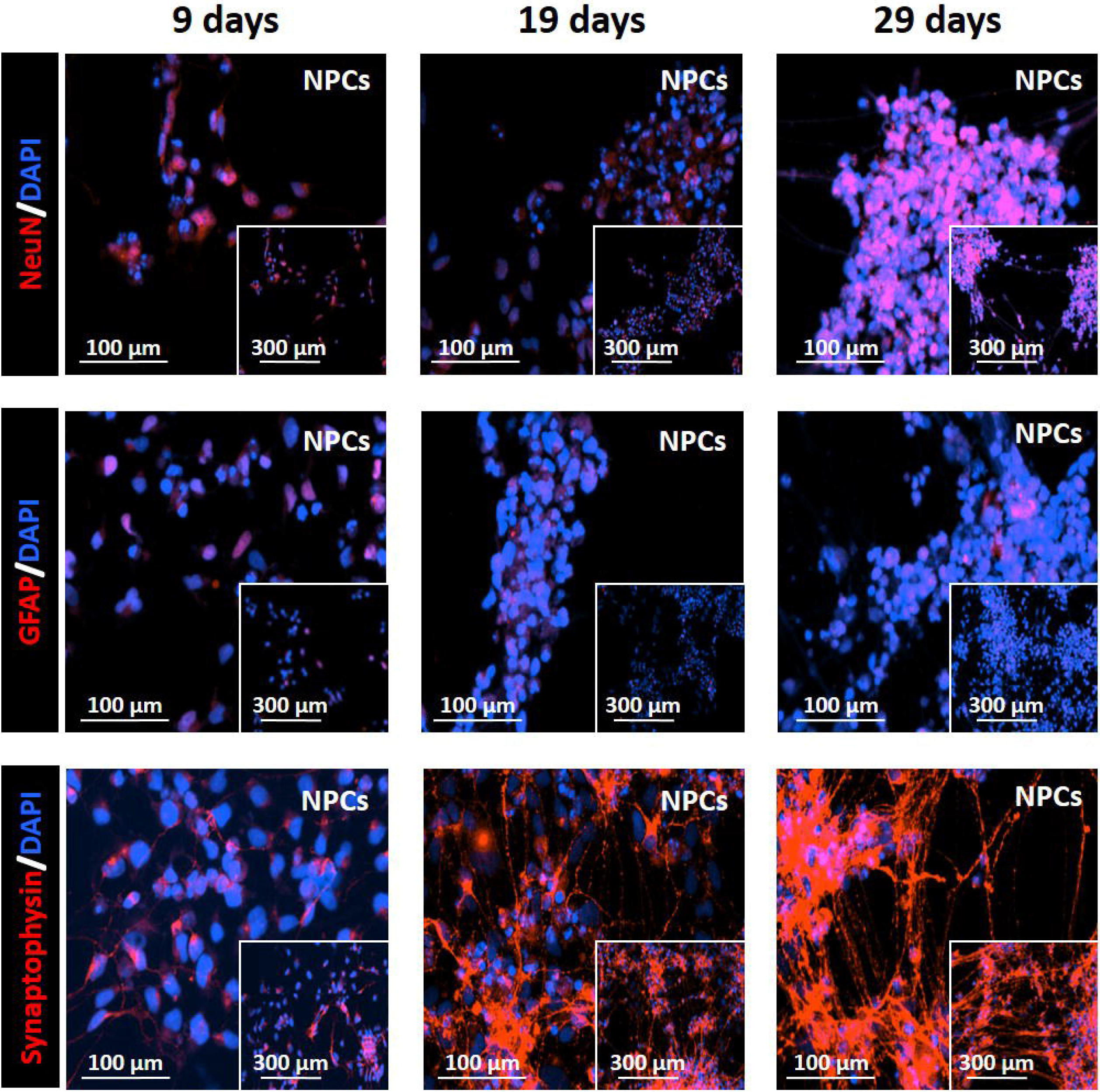
NPCs differentiation immunofluorescence staining. Representative images showing NeuN, GFAP and Synaptophysin protein expression at 9, 19 and 29 days of NPCs incubation in differentiation medium (20 x magnification, scale bar 100 μm, with 20 x magnification 4×4 montage inset, scale bar 300 μm).

To confirm the above observations, we explored if gene expression profiles of different transcription factors and molecular markers changed in our *in vitro* model across consecutive stages of driven neuronal differentiation. For that purpose, we measured mRNA expression levels of Neuronal Differentiation 1 *(NEUROD1)*, Neural Cell Adhesion Molecule 1 *(NCAM1)*, Tubulin Beta 3 Class III *(TUBB3)*, RNA Binding Fox-1 Homolog 3 *(RBFOX3)*, Calbindin 1 *(CALB1)* and *GFAP* genes by RT-qPCR, regardless of the treatment with Aβ (Figure 3).

**Figure 3.**
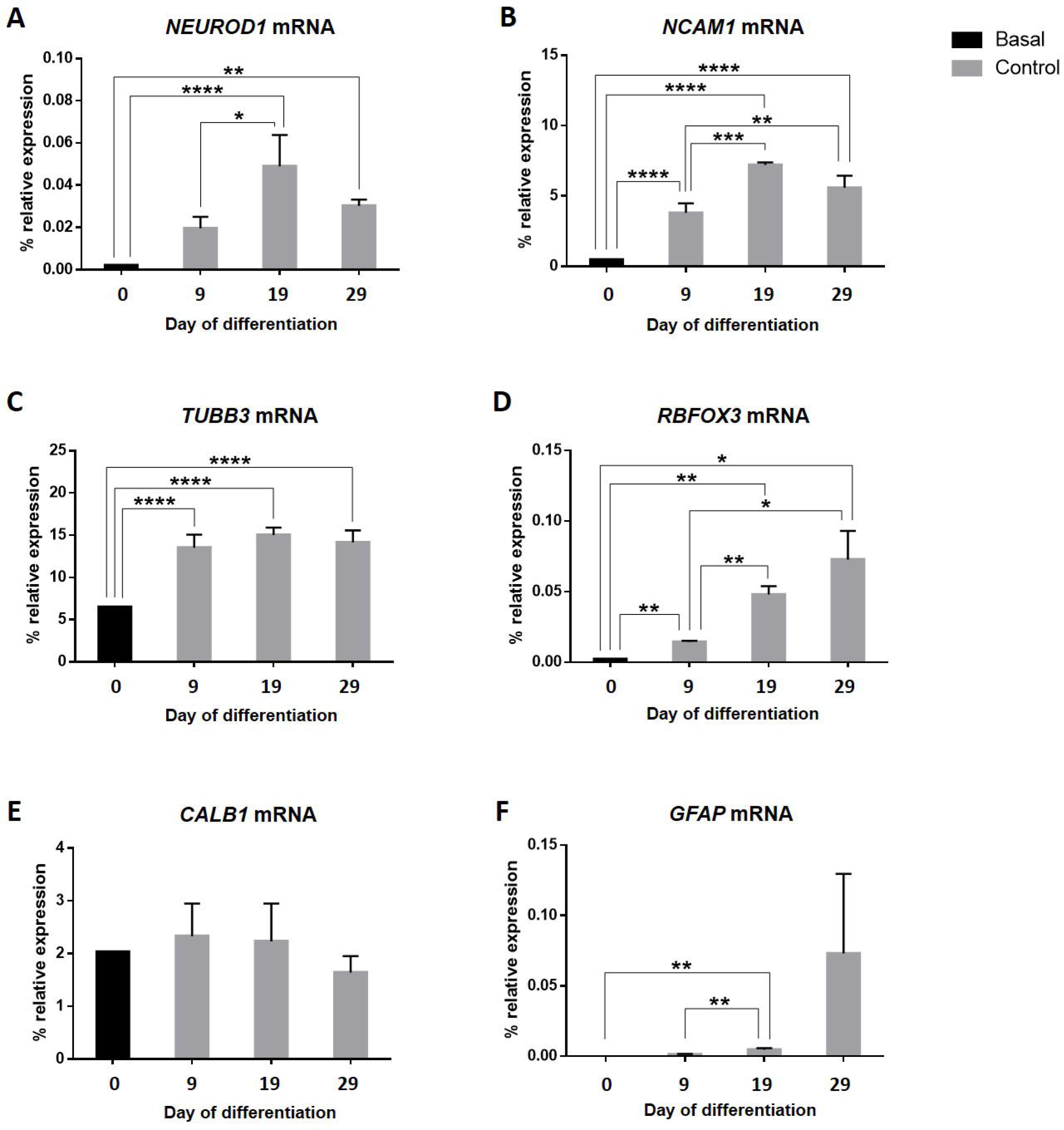
Gene expression profiles of analyzed genes *NEUROD1* (A), *NCAM1* (B), *TUBB3* (C), *RBFOX3* (D), *CALB1* (3) and *GFAP* (F). Bar graphs show the mRNA percentage of relative expression of each gene relative to the geometric mean of *ACTB* and *GAPDH* housekeeping genes expression for basal or control NPCs at each time point of culture. Vertical lines represent the standard error of the mean (SEM). *p-value < 0.05; **p-value < 0.01; ***p-value < 0.001; ****p-value < 0.0001.

*NEUROD1* mRNA expression levels of NPCs cultured in differentiation medium increased from 9 to 19 days [mean difference±standard error (SE), 0.03±0.009; p-value=0.048]. Moreover, we quantified a statistically significant elevation of mRNA expression for this basic helix-loop-helix (bHLH) TF at 19 (0.047±0.008; p-value<0.0001) and 29 days (0.027±0.008; p-value=0.009) compared with basal cells.

In our *in vitro* model, *NCAM1* mRNA expression overlapped that of *NEUROD1* gene, although we found a statistically significant raise since the addition of differentiation medium to the cell culture (ANOVA, p-value<0.001), which is more pronounced at 19 days (5.956±0.468); p-value<0.0001). Our results also displayed statistically significant differences between 9 to 19 days (3.122±0.599; p-value<0.001), 9 to 29 days (2.554±0.599; p-value=0.002) and between basal cells and any of the other time points: from 0 to 9 days (2.834±0.468; p-value<0.0001) and from 0 to 29 days (5.389±0.468; p-value<0.0001).

Once the proliferation medium was exchanged by differentiation medium, NPCs began to express *TUBB3* mRNA, a gene marker that plays a critical role in proper axon guidance and maintenance. This increase remained constant over time comparing with basal cells (6.288±1.002 from 0 to 9 days, 7.351±1.002 from 0 to 19 days and 7.412±1.002 from 0 to 29 days; with a p-value<0.0001 for all of them). However, no changes were observed between first, second and third time points.

*RBFOX3* encodes the NeuN antigen that has been widely used as a marker for post-mitotic neurons. In our study, RBFOX3 mRNA expression grows in time proving the successful achievement of progenitor-to-neuron differentiation. This progressive increase showed statistically significant differences in mRNA expression between 0 and 9 days (0.0105±0.001; p-value=0.001), 9 and 19 days (0.03±0.005; p-value=0.007) or 9 and 29 days (0.057±0.014; p-value=0.047). Besides, all other differences between any time point with respect to basal cells were also found statistically significant: from 0 to 19 days (0.04±0.005; p-value=0.002) and from 0 to 29 days (0.067±0.014; p-value=0.024).

In relation to *CALB1* mRNA expression, and given that this gene encodes a protein expressed in mature granule cells, we did not detect any significant change.

*GFAP* mRNA expression observed coincided in time with that of the neuron lineage-specific gene marker *RBFOX3*, and it was not present in the proliferative stage, at time zero. On the contrary, we noticed a slow raise of *GFAP* expression at 19 days, which was statistically significant compared with basal cells (0.004±0.001; p-value=0.004) and 9 days of differentiation (0.003±0.001; p-value=0.009). This pointed to the existence of NPCs derived astrocytes in the culture.

None of the studied genes showed significant differences of mRNA expression between 19 days and the end time point.

### Effect of Aβ peptide addition on cultured NPCs during neurogenesis stages

In order to mimic the AD context, we exposed NPCs to Aβ peptide once a week throughout the incubation period with differentiation medium. Thus, we wanted to assess whether expression levels of the genes chosen to characterize the stage of neurogenesis in culture were altered due to Aβ peptide addition.

We found transient treatment-specific differences in mRNA expression for some of the studied lineage-specific genes (Figure 4). Aβ peptide reduced *NCAM1* expression [mean of differences±standard error of the mean (SEM), 1.572±0.343]; p-value=0.044) at 19 days and *TUBB3* (1.518±0.296) and *RBFOX3* (0.004±0.0003); p-value=0.008) expression at 9 days. Interestingly, such decreases occurred at the beginning or in between the studied time window, but these differences were no longer significant at the end time point.

**Figure 4.**
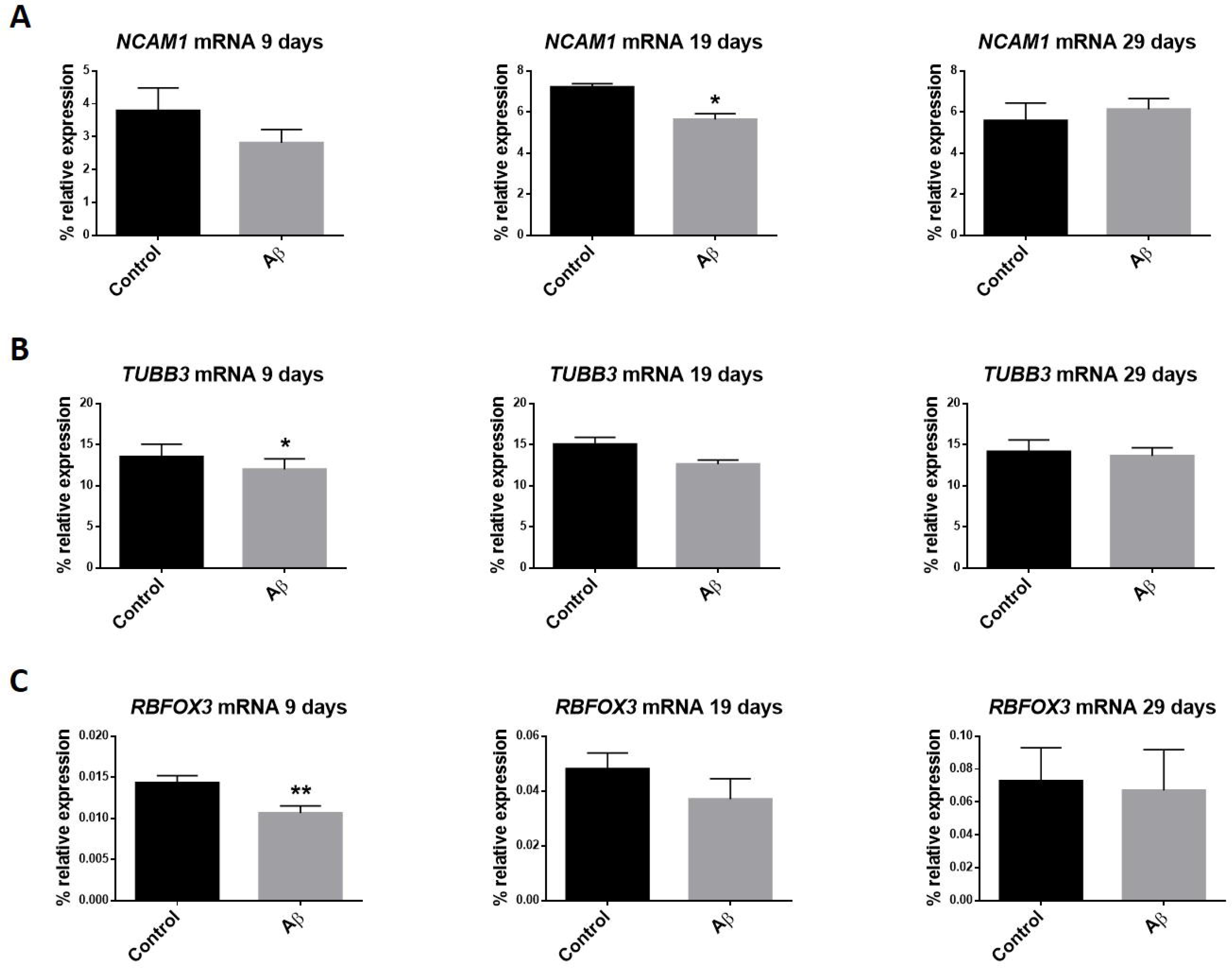
Aβ administration effect during the differentiation process time course. mRNA expression of *NCAM1* (A), *TUBB3* (B) and *RBFOX3* (C) genes relative to the geometric mean of *ACTB* and *GAPDH* housekeeping genes expression was determined for control and Aβ treated NPCs. Vertical lines represent the SEM. *p value < 0.05; **p value < 0.01; ***p value < 0.001; ****p value < 0.0001.

## DISCUSSION

Nowadays, a comprehensive overview has emerged about the stages of AHN development. This complex multistep process can be divided into four phases: a precursor cell phase, an early survival phase, a postmitotic maturation phase, and a late survival phase. Type 1 radial glia-like cells (RGLs) represent the NSC population and can give rise to TAPs (type 2 cells), with first glial (type 2a) and then neuronal (type 2b) phenotype. Through a migratory neuroblast-like stage (type 3), lineage-committed cells exit the cell cycle ahead of maturation into dentate granule neurons functionally integrated into the hippocampal circuitry (Kempermann, Song and Gage, 2015; Zhang and Jiao, 2015). Based on cell morphology, TFs expression and a set of marker proteins, distinct milestones have been established (Kempermann, Song and Gage, 2015). In this study, we examined the expression dynamics of key markers to characterize a directed human NPCs differentiation model across different developmental stages (Figure 5).

**Figure 5.**
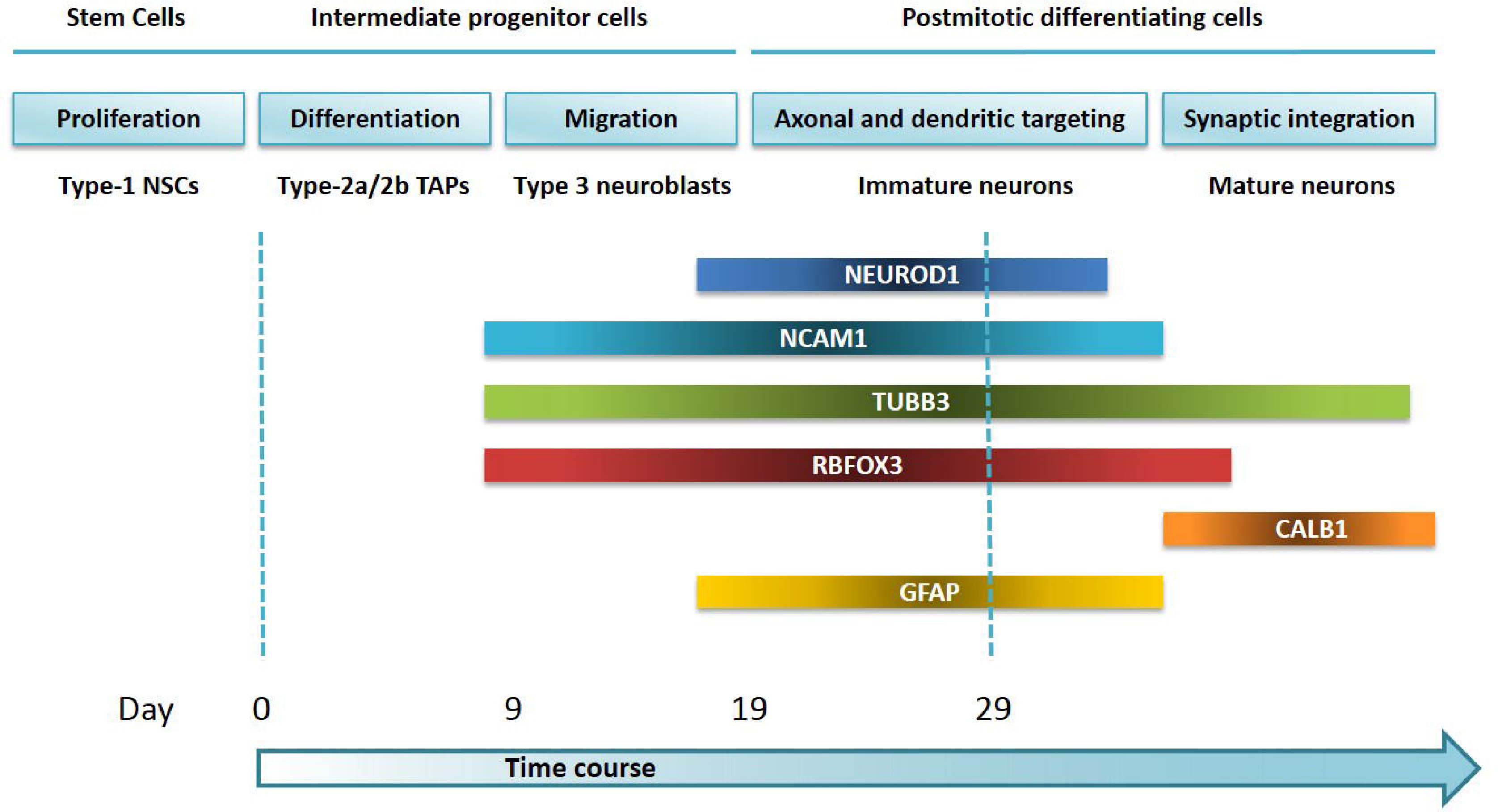
Expression pattern of AHN lineage specific genes studied. The diagram illustrates *NEUROD1, NCAM1, TUBB3, RBFOX3, CALB1* and *GFAP* expression profiles during the directed neuronal differentiation of our time window NPCs culture model, according to the developmental stages of AHN within the neurogenic niche of the DG.

During stage 1 (proliferation phase), newly generated cells express GFAP. However, we do not detect differences in *GFAP* expression until day 19 after the addition of differentiation medium. This leads to believe that our *in vitro* NPCs culture window starts after the proliferative phase, since type-2 cells (differentiation phase) lose the GFAP marker (Zhang and Jiao, 2015). Then, knowing that, in contrast to their *in vivo* counterparts in SGZ of the brain, the *in vitro* expanded NSCs are less neurogenic and mainly biased towards an astrocytic fate upon differentiation (Azari, 2013), *GFAP* expression at day 19 would correspond to a subset of astrocytes present in our NPCs culture (Pierret *et al.*, 2010).

In stage 3 (migration phase) migrating neuroblasts display the polysialylated form of NCAM (PSA-NCAM), a marker that arrives at the late strategy of adult neurogenesis and seems to persist in young postmitotic neurons (von Bohlen Und Halbach, 2007). Accordingly, our results suggest the existence of a plateau between 19 and 29 days for *NCAM1* mRNA expression. Most PSA-NCAM-positive cells express NeuroD and NeuN, but not GFAP, which argues in our favor with respect to our above mentioned findings (von Bohlen Und Halbach, 2007). bHLH TF *NEUROD1*, expressed in later stages of neuronal commitment, plays an essential role in the differentiation and survival of neuronal precursors. NeuroD1 deletion leads to new granule neurons depletion and their failure to integrate in the DG (Shohayeb *et al.*, 2018). In line with this, we observed a rise of *NEUROD1* expression during our culture time window, as reported by Xuan Yu et al. (Yu *et al.*, 2014). Meanwhile, NeuroD expression can also be detected in PSA-NCAM-positive cells, precedes it (von Bohlen Und Halbach, 2007) and reaches the highest point in late stage type 2b and type 3 cells (Hsieh, 2012), although we did not identify this peak in our model. On its part, once the newly generated neurons become postmitotic, they begin to express NeuN marker, which is consistent with an earlier *RBFOX3* mRNA expression in our model. We found how *RBFOX3* expression increased until 19-29 days of differentiation, showing a similar expression profile than *NCAM1*.

Next, cells become postmitotic entering stage 4 (axonal and dendritic targeting). Immature neurons still express PSA-NCAM and, at the same time, can also be marked by NeuN. Also involved in axon guidance and maintenance, *TUBB3* arises, encoding a class III member of the beta tubulin protein family, characteristic of early postmitotic and differentiated neurons and some mitotically active neuronal precursors. This is consistent with the increase of *TUBB3* mRNA detected in our model, prior to be translated to protein. *TUBB3* mRNA expression persists in neurons displaying high complexity and electrophysiological properties, which show immunoreactivity for NeuN and thus represent postmitotic neurons (von Bohlen Und Halbach, 2007).

Finally, mature granule cells establish their synaptic contacts and become functionally integrated in the hippocampus in stage 5 (synaptic integration), expressing calbindin together with NeuN but not co-expressing PSA-NCAM (von Bohlen Und Halbach, 2007). Our group did not find variations in *CALB1* mRNA expression within the analyzed culture time window that might be supposedly expected later in time.

Therefore, based on NPCs monolayer cultured with a medium that accelerates neuronal differentiation by enhancing synaptic activity (Satir *et al.*, 2020), we achieved the challenge of developing a less time consuming differentiation strategy that approximates the *in vivo* developmental program of human hippocampal DG, which differs from that of the SVZ (Ertaylan *et al.*, 2014), as we were able to generate neurons potentially expressing many features of the process of AHN.

In the AD context, it is known that Aβ peptides are generated after cleavage of APP by y-secretase in the amyloidogenic pathway (Coronel *et al.*, 2018). Physiological concentration of Aβ peptides in the brain revealed a positive effect on neuroplasticity and learning, showing improved hippocampal long-term potentiation (LTP), while high nanomolar Aβ administration resulted in impaired cognition (Lazarevic *et al.*, 2017; Garcia-Osta and Alberini, 2009), what reflects an hormetic nature (Puzzo, Privitera and Palmeri, 2012). Because extracellular concentrations of Aβ in normal brain have been estimated to low picomolar levels, in our experiments, we chose a concentration of Aβ peptide 1-42 in the nanomolar range (50 nM) to be added once a week during 29 days of culture, a single dose determined on the average of those used by Gulisano et al. (Gulisano *et al.*, 2018) and Malmsten et al. (Malmsten *et al.*, 2014b)

It has been reported that oligomeric synthetic Aβ peptide 1-42 decreases human NSC proliferative potential and appears to favor glial differentiation, reducing neuronal cell fates (Coronel *et al.*, 2018), or suppresses the number of functional human ES cells-derived neurons (Wicklund *et al.*, 2010). Instead, Bernabeu-Zornoza et al. showed that 1 μM monomeric Aβ peptide 1-42 promoted human NSCs proliferation by increasing the glial precursors pool, without affecting neurogenesis (Coronel *et al.*, 2019). And, on the other hand, differentiating neurospheres exposed to fibrillar Aβ decreased neuronal differentiation and induced gliogenesis (Wicklund *et al.*, 2010; Malmsten *et al.*, 2014a). Controversies existing in the field might be due to Aβ isoforms, peptides concentration, aggregation state, administration times or type of NSCs/NPCs from different species or culture systems used in each experiment (Coronel *et al.*, 2019).

In our case, some of the genes we analyzed showed a decrease in mRNA expression with our Aβ treatment scheme. This transient effect was evident at 9 or 19 days but disappeared at 29 days. It suggests that, despite affecting genes involved in neurogenesis fate, probably before cells maturation, and causing a decrease in differentiation, nanomolar range concentration Aβ addition is somehow counteracted in the long term. A time-dependent reversal in the effects of picomolar Aβ on synaptic plasticity and memory was already seen by Koppensteiner et al., attributable to the enzyme neprilysin, whose levels are reduced with aging and in AD patients brains (Koppensteiner *et al.*, 2016). Strikingly, a study in which mutant APP was overexpressed to ensure Aβ release exclusively by mature neurons found neither a positive nor a negative effect in AHN (Tincer *et al.*, 2016). Hence, our simplistic model might shed light about early AD neurogenesis events, before Aβ deposition cannot be overcome.

A transcriptomic analysis of several human AD profiles demonstrated upregulation of neural progenitor markers expression and downregulation of later neurogenic markers, hypothesizing that neurogenesis is reduced in AD due to compromised maturation (Gatt *et al.*, 2019). Interestingly, they showed downregulation of *NCAM1* expression in early AD hippocampus and also of *NCAM1, TUBB* and *RBFOX3* in late AD hippocampus, which is precisely in line with our Aβ culture treatment results. And recently, Moreno-Jimenez et al. provided evidence for substantial impairment maturation underlying AD progression. Importantly, they identified a decline in doublecortin-expressing cells that co-expressed PSA-NCAM in the DG starting at Braak stage III, followed by a reduction in the expression of NeuN and βIII-tubulin, among others, at some of the subsequent stages of the disease (Moreno-Jimenez *et al.*, 2019).

AHN confers a unique mode of plasticity to the mature mammalian brain. Research about it requires non-invasive monitoring to understand its life-long impact (Bond, Ming and Song, 2015). Easier than manipulating NSCs, in part because of the time-saving, our NPCs model facilitates to study gene expression levels from an *in vitro* cell culture platform that can generate multiple neuronal and glial types in a human genetic context (Efthymiou *et al.*, 2014). Moreover, this straightforward approach helps us to recapitulate AD to deepen our understanding of the alterations affecting specific lineage cell types, even to observe early pathological changes, possibly associated with the prodromal phase of the disease. Nonetheless, SGZ NSCs generate dentate granule neurons and astrocytes while, once propagated in culture with the adequate concentrations of growth factors, they produce all three neural lineages (Bond, Ming and Song, 2015). On the other hand, other cell types are involved in pathogenesis, especially microglia that play a major role, together with neuroinflammation, in AD risk and progression. In consequence, it should be noted that the high potential of NSCs/NPCs could be limited by the characteristics of the *in vivo* niche environment.

Finally, the development of AHN monitoring methods as a biomarker of cognitive function in live individuals will be crucial to stage AD progress. Moreover, to exploit the utility of transcription factor reprogramming to preserve endogenous AHN might contribute to cognitive resilience in AD (Gatt *et al.*, 2019). However, despite exciting, the prospect of using adult NSCs therapeutically as a regenerative source needs to address neuronal integration and its impact into host mature neural circuits (Bond, Ming and Song, 2015). It will involve strategies to accomplish the NSC pool maintenance, generation of correct neuronal subtypes, suppression of glial fates, and differentiation and survival of immature neurons (Hsieh, 2012).

## CONCLUSIONS

All things considered, this work provides a transcriptional profile of a number of genes involved in particular stages of the AHN process to get a detailed understanding of the lineage restricted fate during human neuronal differentiation. Moreover, by the administration of Aβ peptide 1-42 to our human NPCs culture model, we observed similar findings to those obtained in human AD samples relative to *NCAM1, TUBB3* and *RBFOX3* genes expression, possibly offering an *in vitro* opportunity to study AHN impairment in the AD context.

## SUPPLEMENTARY MATERIALS

Supplementary Table S1. RT-qPCR primers.

Supplementary Figure S1. Ki67 protein expression.

## AUTHOR CONTRIBUTIONS

IBL contributed to study concept and design, running experiments, analysis and interpretation of data, figure designing and drawing and drafting/revising the manuscript for content. JC contributed to design and revising manuscript for content. AU contributed to figure designing and drawing and revising the manuscript for content. BA contributed to statistical analysis and revising the manuscript for content. EMGO contributed to design and revising manuscript for content. MR contributed to running experiments. DRPR contributed to interpretation of data and revising manuscript for content. MM contributed to study concept and design, analysis and interpretation of data, statistical analysis, study supervision, drafting/revising the manuscript for content and obtaining funding.

## FUNDING

This research was funded by the Spanish Government through a grant from the Institute of Health Carlos III (FIS PI17/02218), jointly funded by European Regional Development Fund (ERDF), European Union, “A way of shaping Europe”; the Trans-Pyrenean Biomedical Research Network (REFBIO II-MOMENEU project) and Government of Navarra through two grants from Department of Industry of Government of Navarra (“PI058 iBEAS-Plus” and “PI055 iBEAS-Plus”). In addition, AUC received a grant “Doctorandos industriales 2018-2020” and Predoctoral grant (2019) founded by Department of Industry and Health of Government of Navarra. MM received a grant “Programa de intensificación” founded by Fundación Bancaria “la Caixa” and Fundación Caja-Navarra and “Contrato de intensificación” from the Institute of Health Carlos III (INT19/00029).

## ACKNOWLEDGMENTS

We want to kindly thank Valle Coca (Navarrabiomed BrainBank, technical support), Paula Aldaz Ph.D and Imanol Arozarena Ph.D (Cancer Signalling Research Unit, Navarrabiomed, technical and scientific support), Natalia Ramirez Ph.D (Haematological Oncology Research Unit, Navarrabiomed, scientific support), Ibai Tamayo Ph.D, Arkaitz Galbete Ph.D and Julián Librero Ph.D (Methodology Unit, Navarrabiomed, technical support) for their help.

## COMPETING INTERESTS

The authors declare that they have no conflict of interest.

## ABBREVIATIONS

Aβ: amyloid β
AD: Alzheimer’s disease
AHN: adult hippocampal neurogenesis
AN OVA: twoway analysis of variance
APP: amyloid precursor protein
bHLH: basic helix-loop-helix
*CALB1*: calbindin 1
cDNA: complementary DNA
DG: dentate gyrus
ES: embryonic stem
*GFAP*: glial fibrillary acidic protein
HSD: honestly significant difference
IF: immunofluorescence
iPSCs: induced pluripotent stem cells
NCAM1: Neural Cell Adhesion Molecule 1
NeuN: neuronal nuclei
NEUROD1: Neuronal Differentiation 1
NPCs: neural progenitor cells
NSCs: neural stem cells
PSA-NCAM: polysialylated form of NCAM
PSCs: pluripotent stem cells
RBFOX3: RNA Binding Fox-1 Homolog 3
RGLs: radial glia-like cells
RT-qPCR: real time quantitative PCR
SE: standard error
SEM: standard error of the mean
SGZ: subgranular zone
SVZ: subventricular zone
TAPs: transient amplifying progenitors
TFs: transcription factors
TUBB3: Tubulin Beta 3 Class III.

